# Automated, predictive, and interpretable inference of *C. elegans* escape dynamics

**DOI:** 10.1101/426387

**Authors:** Bryan C. Daniels, William S. Ryu, Ilya Nemenman

## Abstract

The roundworm *C. elegans* exhibits robust escape behavior in response to rapidly rising temperature. The behavior lasts for a few seconds, shows history dependence, involves both sensory and motor systems, and is too complicated to model mechanistically using currently available knowledge. Instead we model the process phenomenologically, and we use the *Sir Isaac* dynamical inference platform to infer the model in a fully automated fashion directly from experimental data. The inferred model requires incorporation of an unobserved dynamical variable, and is biologically interpretable. The model makes accurate predictions about the dynamics of the worm behavior, and it can be used to characterize the functional logic of the dynamical system underlying the escape response. This work illustrates the power of modern artificial intelligence to aid in discovery of accurate and interpretable models of complex natural systems.

---

The quantitative biology revolution of the recent decades has resulted in an unprecedented ability to measure dynamics of complex biological systems in response to perturbations with the accuracy previously reserved for inanimate, physical systems. For example, the entire escape behavior of a roundworm *Caenorhabditis elegans* in response to a noxious temperature stimulus can be measured for many seconds in hundreds of worms [1, 2]. At the same time, theoretical understanding of such living dynamical systems has lagged behind, largely because, in the absence of symmetries, averaging, and small parameters to guide our intuition, building mathematical models of such complex biological processes has remained a very delicate art. Recent years have seen emergence of *automated modeling* approaches, which use modern machine learning methods to automatically infer the dynamical laws underlying a studied experimental system and predict its future dynamics [3–14]. However, arguably, these methods have not yet been applied to *any* real experimental data with dynamics of *a priori* unknown structure to produce interpretable dynamical representations of the system. Thus their ability to build not just statistical but *physical* models of data [15], which are interpretable by a human, answer interesting scientific questions, and guide future discovery, remains unclear.

Here we apply the *Sir Isaac* platform for automated inference of dynamical equations underlying time series data to infer a model of the *C. elegans* escape response, averaged over a population of worms. We show that *Sir Isaac* is able to fit not only the observed data, but also to make predictions about the worm dynamics that extend beyond the data used for training. The inferred optimal model is fully interpretable, with the identified interactions and the inferred latent dynamical variable being biologically meaningful. And by analysing the dynamical structure of the model—number of dynamic variables, number of attractors (distinct behaviors), etc.—we can generalize these results across many biophysical systems.

## Results

### Automated Dynamical Inference

*Sir Isaac* [7, 8] is one of the new generation of machine learning algorithms able to infer a dynamical model of time series data, with the model expressed in terms of a system of differential equations. Compared to other approaches, *Sir Isaac* is able to infer dynamics (at least for synthetic test systems) that are (i) relatively low-dimensional, (ii) have unobserved (hidden or latent) variables, (iii) have arbitrary nonlinearities, (iv) rely only on noisy measurements of the system’s state variables, and not of the rate of change of these variables, and (v) are expressed in terms of an interpretable system of coupled differential equations. Briefly, the algorithm sets up a complete and nested hierarchy of nonlinear dynamical models. *Nestedness* means that each next model in the hierarchy is more complex (in the sense of having a larger explanatory power) [16–18] than the previous one, and includes it as a special case. *Completeness* means that *any* sufficiently general dynamics can be approximated arbitrarily well by some model within the hierarchy. Two such hierarchies have been developed, one based on S-systems [19] and the other on sigmoidal networks [20]. Both progressively add hidden dynamical variables to the model, and then couple them to the previously introduced variables using nonlinear interactions of specific forms. *Sir Isaac* then uses a semi-analytical formulation of Bayesian model selection [8, 18, 21, 22] to choose the model in the hierarchy that best balances the quality of fit versus overfitting and is, therefore, expected to produce the best generalization. The sigmoidal network hierarchy is especially well-suited to modeling biological systems, where rates of change of variables usually show saturation, and it will be the sole focus of our study.

### Experimental model system

Nociception evokes a rapid escape behavior designed to protect the animal from potential harm [23, 24]. *C. elegans*, a small nematode with a simple nervous system, is a classic model organism used in the study of nociception. A variety of studies have used *C. elegans* to elucidate genes and neurons mediating nociception to a variety of aversive stimuli including high osmolarity and mechanical, chemical, and thermal stimuli [25–28]. However, a complete dynamic understanding of the escape response at the neuronal, let alone the molecular, level is not fully known. Recent studies have quantified the behavioral escape response of the worm when thermally stimulated with laser heating [1, 2], and these data will be the focus of our study. The response is dynamic: when the stimulus is applied to the animal’s head, it quickly withdraws, briefly accelerating backwards, and eventually returns to forward motion, usually in a different direction. Various features of this response change with the level of laser heating, such as the length of time moving in the reverse direction and the maximum speed attained.

### Fits and Predictions

We use the worm center-of-mass speed, *v*, as the variable whose dynamics needs to be explained in response to the laser heating pulse. We define *v* > 0 as the worm crawling forward and *v* < 0 as the worm retreating backwards. The input to the model is the underlying temperature, *h*(*t*), which can be approximated as *h*(*t*) = *Ih*_0_(*t*), where *I* is the experimentally controlled laser current, and *h*_0_ is the temperature template, described in *Materials and Methods*. Based on trajectories of 201 worms in response to laser currents ranging between 9.6 and 177.4 mA, we let *Sir Isaac* determine the most likely dynamical system explaining this data within the sigmoidal networks model class [8] (see *Materials and Methods* for a detailed description of the modeling and inference). The inferred model has a latent (unobserved) dynamical variable, hereafter referred to as *x*_2_, in addition to the speed. *v* and *x*_2_ are coupled by nonlinear interactions. However, some of these nonlinear interactions may be insignificant, and may be present simply because the nested hierarchy introduces them before some other interaction terms that are necessary to explain the data. Thus we reduce the model by setting parameters that are small to zero one by one and in various combinations, refitting such reduced models, and using Bayesian Model Selection to choose between the reduced model and the original *Sir Isaac* inferred model. The resulting model is

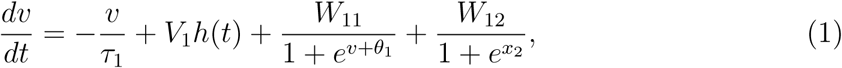

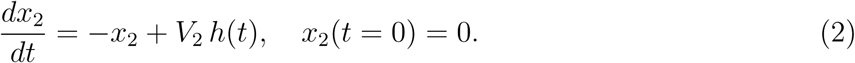

Here *V*_1_, *V*_2_, *W*_11_, *W*_12_ are constants inferred from data, while the model uses the default value of 1.0 s for the characteristic time scale of the dynamics of *x*_2_ (see *Materials and Methods* for the values of the parameters). Interestingly, the inferred model reveals that the latent dynamical variable *x*_2_ is a linear low-pass filtered (integrated) version of the heat signal.

The fits produced by this model are compared to data in Fig. 1(A), showing an excellent agreement (see *Materials and Methods* for quantification of the quality of fits). Surprisingly, the quality of the fit for this automatically generated model is *better* than that of a manually curated model [2]: only about 10% of explainable variance in the data remains unexplained by the model for times between 100 ms and 2 s after the stimulus, compared to about 20% for the manual model, cf. Fig. 5(A).

**Fig. 1.**
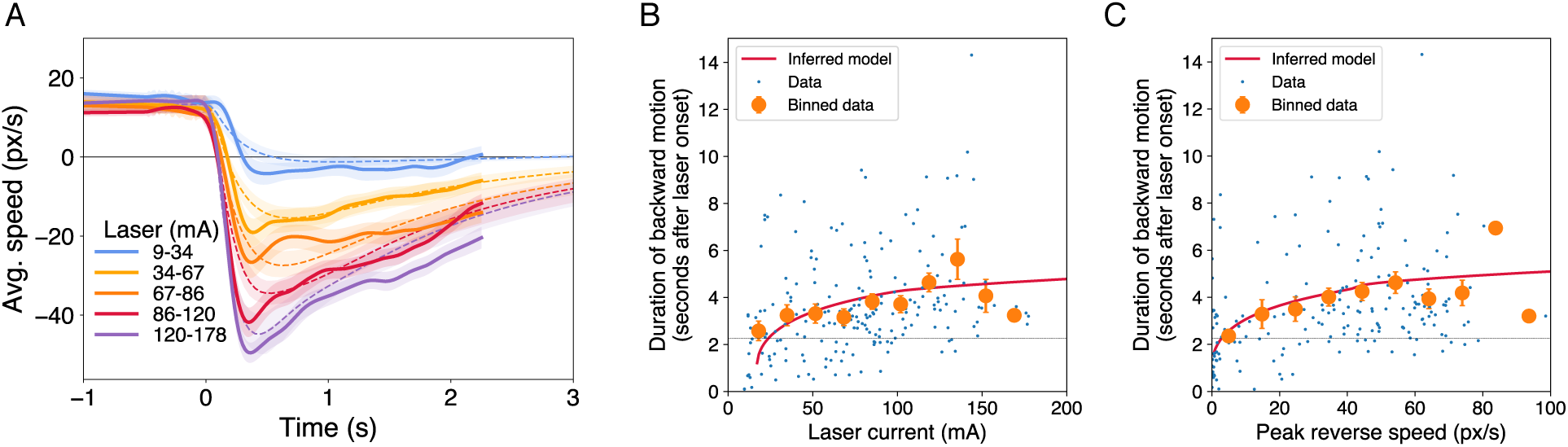
The escape response behavior is fitted and predicted well by the inferred model. (A) Colored lines and shaded bands represent the empirical mean and the standard deviation of the mean, respectively, of the escape response velocity for five groups of worms stimulated with laser currents in different ranges (38, 42, 39, 41, and 41 subjects in each group). Dashed lines and bands of the corresponding color show means and standard deviations of the mean (see *Materials and methods*) of fits to these empirical data by the chosen model. Only the velocity in the range of time [−1, 2.25] s relative to the time of the onset of the laser stimulus was used for fitting. While most worms are still moving backward during this range of time, the inferred model *predicts*, without any additional free parameters, the time at which a worm’s speed again becomes positive both as a function of (B) applied laser current and (C) peak observed worm reverse speed. These predictions (red curves) agree well with the binned averages (orange points with error bars representing the standard error of the mean if the bin has more than five worms) of individual worms’ behavior (blue dots).

However, the quality of the fit is not surprising in itself since the *Sir Isaac* model hierarchy can fit any dynamics using sufficient data. A utility of a mathematical model is in its ability to make *predictions* about data that were not used in fitting. Thus we use the inferred model to predict when the worm will return to forward motion, which usually happens well after the temporal range used for fitting. Figure 1(B,C) compares these predictions with experiments, showing very good predictions. Such ability to extrapolate beyond the training range is usually an indication that the model captures the underlying physics, and is not purely statistical [15], giving us confidence in using the model for inferences about the worm.

### Model analysis

The algorithm has chosen to include a single latent dynamical variable, which is a linear leaky integrator of the experienced temperature. Having access to both the instantaneous stimulus and its integral over the immediate past allows the worm to estimate the rate of change in the stimulus. This agrees with the observation [1] that both the current temperature, as well as the rate of its increase, are noxious for the worm. From this, one could have guessed, perhaps, that *at least* one latent variable (temperature derivative, or temperature at some previous time) is required to properly model the escape response. However, the fact that *Sir Isaac* inferred this from time series data alone and was able to model the data with *exactly* one hidden variable is surprising.

Figure 2 shows the phase portraits of the inferred dynamical model, Eqs. (1, 2), as well as the dynamics of the speed and the heat stimulus *h*. Crucially, we see that there is only one fixed point in the phase space at any instant of time, and the position of this fixed point is affected by the current laser stimulus value. This suggests that, at least at the level of the population-averaged response, the behavior does not involve switching among alternative behaviors defined dynamically as multiple fixed points or limit cycles (e. g., forward and backward motion) with the switching probability influenced by the stimulus [29], but rather the stimulus controls the direction and the speed of the single dominant crawling state.

**Fig. 2.**
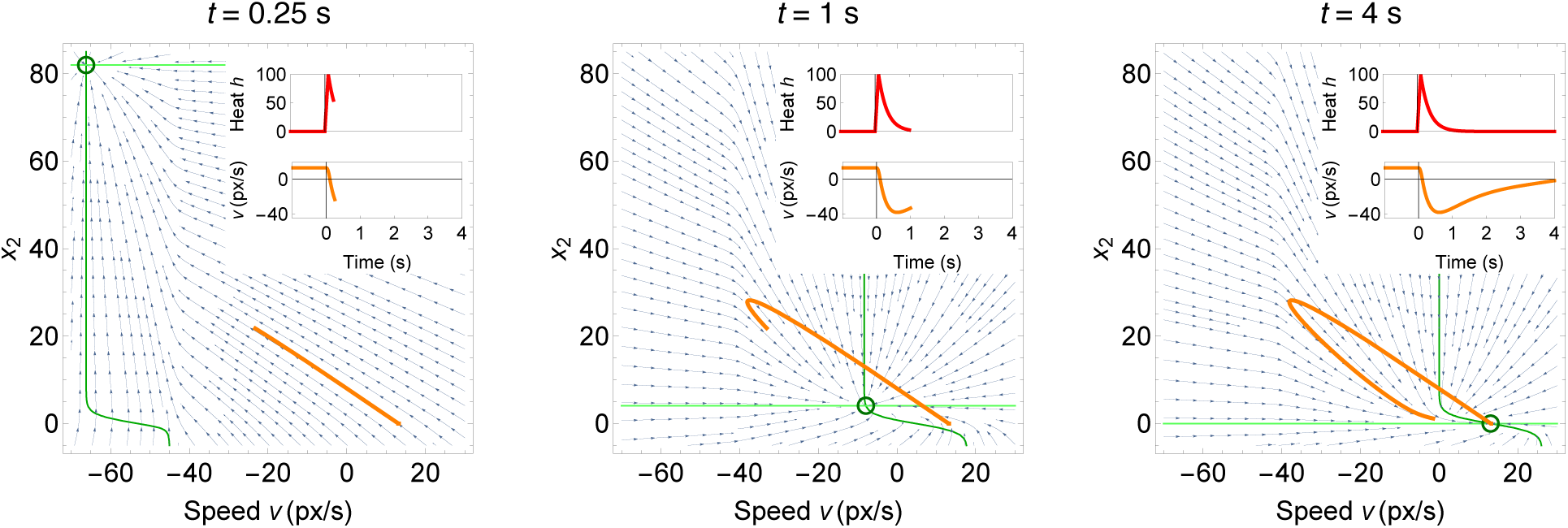
Phase space structure of inferred model. With one hidden variable *x*_2_, the model dynamics can be visualized in the two-dimensional phase space. As the instantaneous heat input *h* returns to zero after a brief pulse (red curve in insets), a single fixed point in the two-dimensional (*v*, *x*_2_) dynamics moves from negative speed (escape) to positive (forward motion). As the velocity of the worm trails the fixed point, this produces first a fast escape and then a slow return to forward motion (speed trajectory in orange in inset and in the phase portrait plots). Blue arrows indicate flow lines, circles indicate stable fixed points, and green lines indicate nullclines (dark green where *dv/dt* = 0 and light green where *dx*_2_*/dt* = 0).

The network diagram of the model in Eqs. (1, 2) is shown in Fig. 3, where we omit the linear degradation terms for *v* and *x*_2_. With the maximum likelihood parameter values, Tbl. III, the model can be interpreted as follows. The hidden variable *x*_2_ is a linear leaky integrator of the heat signal, storing the average recent value of the stimulus over about 1 s. While we do not know which exact neuron can be identified with *x*_2_, the thermosensory neurons AFD, FLP located in head of the worm are strong candidates [30, 31]. The thermosensory neurons AFD respond to changes in temperature and are the primary sensors responsible for thermotaxis [30]. The sensory neuron FLP also is thermosensitive and has a role in the thermal sensory escape response [31]. When *x*_2_ is near zero, the *W*_12_ term is large, and, together with the linear relaxation of the speed, −*v/τ*1, it establishes a constant positive forward motion. We identify this term with the forward drive command interneurons AVB, PVC [32]. After the temperature increases, *x*_2_ grows. It rapidly increases the denominator in the *W*_12_ term and hence shuts down the forward drive. This is again consistent with the literature indicating that the worms pause with even reasonably small temperature perturbations [1]. An additional effect of the stimulus is to directly inject a negative drive −*V*_1_*h* into the dynamics of the velocity. When the stimulus is large, the *V*_1_ term is sufficiently negative to result in the velocity overshooting the pause into the negative, escape range. We identify the *V*_1_ term with the reverse command interneurons AVA, AVD, and AVE, activated by the thermosensory neurons AFD, FLP [32]. When *v* > −*θ*_1_ ≈ −42 px/s, the *W*_11_ term is suppressed. However, during fast escapes this suppression is lifted, activating the positive drive *W*_11_, which leads to faster recovery of the forward velocity. We identify the *W*_11_ term with internal recovery dynamics of the reverse command interneurons whose molecular mechanisms of activity are only partially understood [33]. The velocity does not just relax to zero over some characteristic time, but crosses back into the forward crawl once *x*_2_ has decreased sufficiently to reactivate the *W*_12_ term. Overall, the biological interpretability of the model is striking. And where there is no direct match between known worm biology and the model, the model strongly suggests that we should be looking for specific predicted features, such as the neural and molecular mechanisms for both sensing the heat stimulus and its recent average.

**Fig. 3.**
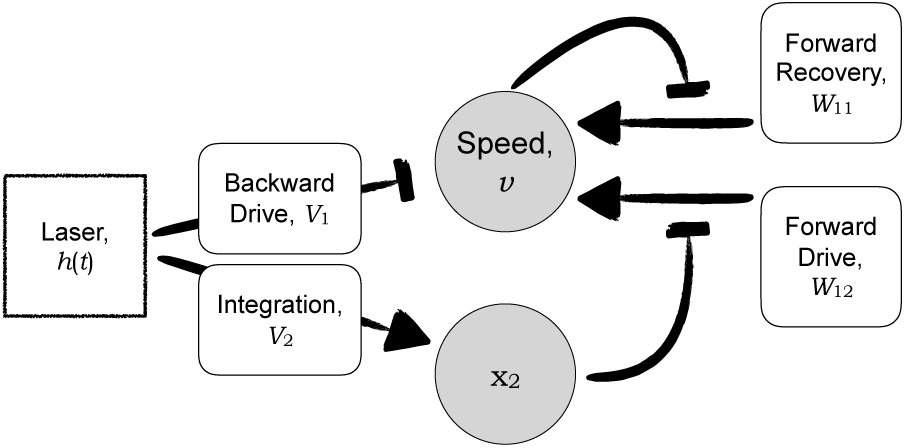
Network diagram of the inferred model. Variables, interactions, parameter values, and biological mechanisms are described in the main text.

Another notable feature of the network diagram is that it is similar to other well-known sensory networks, namely chemotaxis (and the related thermotaxis) in *E. coli* [34, 35] and chemotaxis in *D. discoideum* [36, 37]. In all these cases, the current value of the stimulus is sensed in parallel to the stimulus integrated over the recent time. They are later brought together in a negative feedback loop (*E. coli*) or an incoherent feedforward loop (*D. discoideum*), resulting in various adaptive behaviors. In contrast, the worm’s behavior is more complicated: the ambient temperature participates in a negative feedback loop on the speed through *W*_11_, which results in adaptation. At the same time, the integrated temperature (through *W*_12_) and the current temperature add *coherently* to cause the escape in response to both the temperature and its rate of change. This illustrates the difference between sensing, when an organism needs to respond to stimulus changes only, and escape, where more complicated dynamics is needed.

## Discussion

In this article, we have used modern machine learning to learn the dynamics underlying the temperature escape behavior in *C. elegans*. The resulting automatically inferred model is more accurate than the model curated by hand. It uses the dynamics within what is normally considered as discrete behavioral states to make extremely precise verifiable (and verified) predictions about the behavior of the worm beyond the range of time used for training. The model is fully interpretable, with many of its features having direct biological, mechanistic equivalents. Where such biological equivalents are unknown, the model makes strong predictions of what they should be, and suggests what future experiments need to search for.

One can question if describing the *C. elegans* nociceptive behavior, which typically is viewed as stochastic [1, 2] and switching between discrete states, with the deterministic dynamics approach of *Sir Isaac* is appropriate. The quality of the fits and predictions is an indication that it is. This is likely because (i) the escape is, to a large extent, deterministic, becoming more and more so as the stimulus intensity increases [1]; (ii) on the scale of individual worms, the discretized behavior states (forward, backward, pause, …) have their own internal dynamics, with different time-dependent velocities, which *Sir Isaac* models well, and the boundaries between the states are not highly pronounced; and (iii) our equations model a population of worms, and so even if individual worms were dominated by stochasticity, *Sir Isaac* would do just fine modeling the dynamics of the mean behavior.

Crucially, the model discovers that the behavior, at least of an average worm, is not a simple one-to-one mapping of the input signal: the instantaneous stimulus and its temporally integrated history (one latent variable) are both important for driving the behavior. The behavior is driven by one fixed point in the velocity-memory phase space, and the worm changes its speed while chasing this fixed point, which in turn changes in response to the stimulus. This is in contrast to other possibilities, such as the worm being able to exhibit both the forward and the backward motion at any stimulus value, and the stimulus and its history merely affect the probability of engaging in either of these two behaviors.

This is the first successful application of automatic phenomenological inference to modeling dynamics of complex biological systems, without using the helping constraints imposed by (partial) knowledge of the underlying biology. The emphasis on interpretable, physical models allows extrapolation well beyond the data used for training, which is hard for purely statistical methods. The automation allows for a comprehensive search through the model space, so that the automatically inferred model is better than the human-assembled one, especially when faced with only partial knowledge of the complex system. Even for the best studied biological systems, we do not have the necessary set of measurements to model them from the ground up. Our work illustrates the power of phenomenological modeling approach, which allows for top down modeling, adding interpretable constraints to our understanding of the system.

## Materials and Methods

### Data collection

A detailed description of the experimental methods has been previously published [2]. In summary, we raised wild-type, N2 *C. elegans* using standard methods, incubated at 20°C with food. Individual worms were washed to remove traces of food and placed on the surface of an agar plate for 30 minutes at 20°C to acclimatize. Worms were then transferred to an agar assay plate seeded with bacteria (food) and left to acclimatize for 30 more minutes. The worms were then stimulated with an infra-red laser focused to a diffraction limited spot directed at the “nose” of the worm. The intensity of the stimulus was randomized by selecting a laser current between 0 and 200 mA, with a duration of 0.1s. The temperature increase caused by laser heating was nearly instantaneous and reached up to a maximum of 2°C for our current range. Each worm was stimulated only once and then discarded. Video of each worm’s escape response was recorded at 60 Hz and processed offline using custom programs in LabVIEW and MATLAB.

### Input data

Data used for dynamical inference are as described in Ref. [2]. Speed data for 201 worms were extracted from video frames at 60 Hz and smoothed using a Gaussian kernel of width 500 ms. We use data between 0.5 s before and 2.25 s after the start of the laser stimulus. Aligning the data by laser start time, the stimulus happens at the same time in each trial. Naively used, this can produce models that simply encode a short delay followed by an escape, without requiring the stimulus input. To ensure that instead the stimulus causes the response in the model, for each trial we add a random delay between 0 and 1 s to the time data.

Additionally, we are not interested in capturing any dynamics in the pre-laser free crawling state. If we only use the small amount of pre-laser data measured in this experiment, the inference procedure is free to include models with complicated transient behavior before the stimulus. For this reason, we include a copy of pre-laser data at a fictitious “equilibration time” long before the stimulus time (10 s), artificially forcing the model to develop a pre-stimulus steady state of forward motion. Finally, we weight the pre-laser data such that it appears with equal frequency as post-laser data in fitting, in order that the inference algorithm is not biased toward capturing post-laser behavior more accurately than the pre-laser one.

### Estimating explainable variance

The observed variance in worm speed can be partitioned into that caused by the input (changes in laser current) and that caused by other factors (individual variability, experimental noise, etc.). As in Ref. [2], we focus on our model’s ability to capture the former, “explainable” variance. We treat the latter variance as “unexplainable” by our model, and it is this variance that we use to define uncertainties on datapoints for use in the inference procedure. We estimate these variances by splitting the data into trials with similar laser current *I_µ_* (5 bins), producing variances 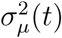 that depend on the laser current bin and on the time relative to laser start. For simplicity, we use a constant uncertainty for all data-points that is an average over laser current bins and times relative to the stimulus onset, which is equal to *σ* = 14.2 px/s. We use this uncertainty when calculating the *χ*^2^ that defines the model’s goodness-of-fit:

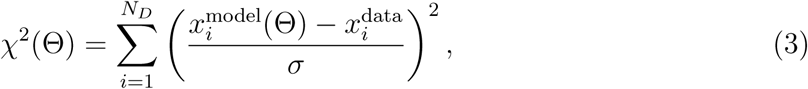

where the index *i* enumerates data points used for the evaluation, *N_D_* is the number of points, and Θ is the vector of all parameters.

### *Sir Isaac* inference algorithm

We use the *Sir Isaac* dynamical inference algorithm [8] to find a set of ODEs that best describes the data without overfitting. Based on previous studies of simulated biological systems, we use the continuous-time sigmoidal network model class, which produces a set of *J* ODEs of the form

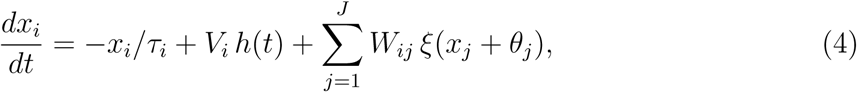

where *ξ*(*y*) = 1/(1 + exp(*y*)) and *h*(*t*) is the sensory input defined in Eq. (7). The algorithm infers both the number of parameters (controlled by the total number of dynamical variables *J*) and the parameter values themselves (the timescales *τ_i_*, interaction strengths *V_i_* and *W_ij_*, and biases *θ_j_*). The first dynamical variable *x*_1_ is taken to be the signed speed of the worm’s center of mass, in pixels per second, with negative values corresponding to backward motion. Further dynamical variables *x_i_* with *i* > 1 correspond to latent (unmeasured) dynamical variables.

The fitting procedure starts by using one datapoint (one random time) from each of a few trials, then gradually adds trials and eventually multiple datapoints per trial, refitting model parameters at each step. The resulting model fits are scored based on their performance in predicting the entire time series (see Fig. 4 for the fit quality). When the performance and model complexity of the winning model saturate, we use the resulting model as our description of the system. In this way, parameters are fit using only a small subset of the available data—we find that using *∼* 2 randomly chosen timepoints per trial (out of the total 165) is sufficient, cf. Fig. 4. This approach significantly reduces computational effort (which scales linearly in the number of datapoints used) and minimizes the effects of correlations between datapoints that are close to one another in time. Finally, it prevents the optimization from getting stuck at local minima and saddles, which change as new data points are added, or data is randomized. These reasons are similar to the reasons behind stochastic gradient descent approaches [38]. The developed software is available from https://github.com/EmoryUniversityTheoreticalBiophysics/SirIsaac.

**Fig. 4.**
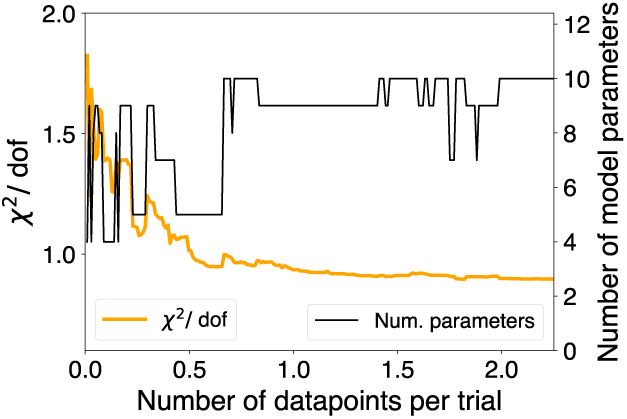
Monitoring goodness of fit in the process of model inference. *Sir Isaac* adds data gradually to aid in parameter fitting. As data is added, the selected model includes more detail (number of model parameters in black) until it saturates to the 10 parameter model we use. The goodness of fit (*χ*^2^ per degree of freedom, in yellow) is measured using all data, including time points not used in model fitting.

### Inferred models

The inference procedure produces the following differential equations:

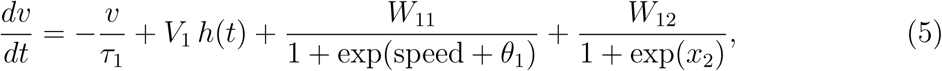

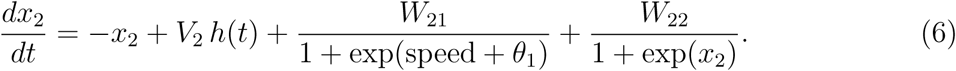

The model includes one latent dynamical variable *x*_2_. The maximum likelihood fit parameters are shown in Table II. (Note that the selected model did not include variables *τ*_2_ and *θ*_2_, so they are set to their default values: *τ*_2_ = 1 and *θ*_2_ = 0.)

**Table I.** Parameters used for input data, dynamical model inference, and the dynamics of the sensory input. The inference parameters quoted are used in the *Sir Isaac* algorithm: “complexity stepsize” sets which models in the model hierarchy are tested, with 1 indicating that no models are skipped as parameters are added to make models more complex; here “ensemble” refers to the parameter ensemble used to avoid local minima during parameter fitting; “avegtol” and “maxiter” are parameters controlling the local minimization phase of parameter fitting; “stopfittingN” sets the number of higher-complexity models needed to be shown to have poorer performance before selecting a given model and finishing the optimization; “connection order” and “type order” set the order of adding parameters within the model hierarchy [7]; and “prior *σ*” is the standard deviation of the Gaussian priors on all parameters.

|  |  |  |  |
| --- | --- | --- | --- |
| **Input data parameters** |  |  |  |
| equilibration time | 10 s |  |  |
| **Inference parameters** |  |  |  |
| complexity stepsize | 1 | tested ensemble members | 10 |
| maxiter | 100 | ensemble temperature | 100 |
| avegtol | $10^{-2}$ | ensemble generation steps | 1000 |
| stopfittingN | 5 | connection order | ‘node’ |
| type order | ‘last’ | prior $\sigma$ | 100 |
| **Sensory input parameters** |  |  |  |
| $\tau_{\text{on}}$ | 0.1 s | $\tau_{\text{decay}}$ | 0.25 s |

**Table II.** Maximum-likelihood parameters for the model inferred by *Sir Isaac*. We do not calculate parameter co-variances since posteriors are sloppy [39] and non-Gaussian around the maximum likelihood. Instead we estimate the standard deviation of the fits themselves, cf. Fig. 1(A). Parameter values are reported to an accuracy of about one digit beyond the least statistically significant one.

|  |  |  |  |
| --- | --- | --- | --- |
| $\tau_1$ | 1.33 s | $\theta_1$ | 42.1 px/s |
| $V_1$ | -2.30 px/(s<sup>2</sup> mA) | $V_2$ | 1.7 1/(s mA) |
| $W_{11}$ | 81.0 px/s<sup>2</sup> | $W_{12}$ | 44.1 px/s<sup>2</sup> |
| $W_{21}$ | -0.1 s<sup>-1</sup> | $W_{22}$ | 5.5 s<sup>-1</sup> |
| $v(t=0)$ | 12.6 px/s | $x_2(t=0)$ | 1.6 |

We notice that some of the inferred parameters are close to zero, and so we check whether the model can be simplified by setting each of them to zero one at a time and in combinations and then measuring the approximate Bayesian model selection posterior likelihood score [8] for the original *Sir Isaac* inferred model and for each of the reduced models. The original model has the log-likelihood of −197.2, and the best model, Eqs. (1, 2), has the highest Bayesian likelihood of −192.9. This becomes our chosen model, maximum likelihood parameters for which can be found in Table III.

**Table III.** Maximum-likelihood parameters for the reduced model.

|  |  |  |  |
| --- | --- | --- | --- |
| $\tau_1$ | 1.33 s | $\theta_1$ | 42.1 px/s |
| $V_1$ | -2.30 px/(s<sup>2</sup> mA) | $V_2$ | 1.5 1/(s mA) |
| $W_{11}$ | 76. px/s<sup>2</sup> | $W_{12}$ | 19.7 px/s<sup>2</sup> |
| $v(t=0)$ | 12.1 px/s | | |

### Bayesian model of uncertainty

To quantify uncertainty in model structure and parameters, we take a Bayesian approach and sample from the posterior distribution over parameters. Assuming independent Gaussian fluctuations in the data used in fits, the posterior is simply exp (−*χ*^2^(Θ)), with *χ*^2^(Θ) from Eq. (3). We use a standard Metropolis Monte Carlo sampler, as implemented in SloppyCell [40], to take 100 samples (each separated by 1000 Monte Carlo steps) used to quantify uncertainty in the fit, cf. Fig 1(A).

### Model of sensory input

Each experimental trial begins with a forward moving worm, which is stimulated with a laser pulse of duration *τ_on_* = 0.1 s starting at time *t* = 0. The worm’s nose experiences a quick local temperature increase *h*(*t*), which we model as a linear increase during the pulse, with slope proportional to the laser current *I*. The time scale of the temperature decay of the heated area to the ambient temperature due to heat diffusion is 0.15 s [41]. However, the stimulation area is broad (220*µ*m, FWHM), and as the worm retreats, its head with the sensory neurons first moves deeper into the heated area, before the temperature eventually decreases. Thus the dynamics of the sensory input is complex and multiscale. However, since each individual behaves differently, and we do do not measure individual head temperatures, we model the *average* sensory stimulus past the heating period as a single exponential decay with the longer time scale of *τ*_decay_ = 0.25 s:

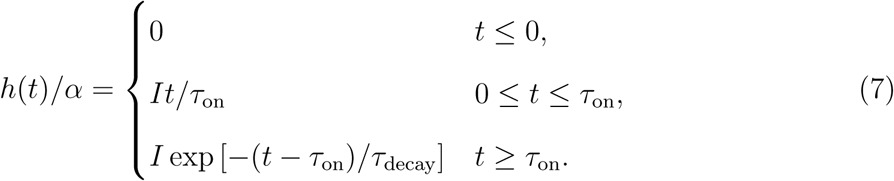

Here *α* has units temperature per unit of laser current. For convenience and without loss of generality, we set *α* = 1 and absorb its definition into the *V_i_* parameters that multiply *h* in Eqs. (5, 6) giving *V_i_* units of the time derivative of *x_i_* per unit current.

### Comparison to the model of Ref. [2]

A quantitative model of *C. elegans* nociceptive response was constructed in Ref. [2]. It involved partitioning worms into actively escaping and (nearly) pausing after a laser stimulation, estimating the probability of pausing as a function of the applied laser current, and finally noticing that the mean response of the active worms is nearly stereotypical, with the response amplitude depending nonlinearly on the stimulus intensity. The same stimulus causes varied response trajectories in individual worms, and this variability is unexplainable in any model that only relates the stimulus to the population averaged response velocity. Of the variability in the response that is explainable by the stimulus, the model of Ref. [2] could not account for ∼ 20% in the range from a few hundred ms to about 2 s post-stimulation. Figure 5(A) shows that *Sir Isaac* our selected model captures *more* of the data variance, leaving only 10% of the explainable variance unexplained over much of the same time range. Figure 5(B,C) illustrates that *Sir Isaac* has recovered the approximate stereotypy in the response to the same accuracy as it is present in the data.

**Fig. 5.**
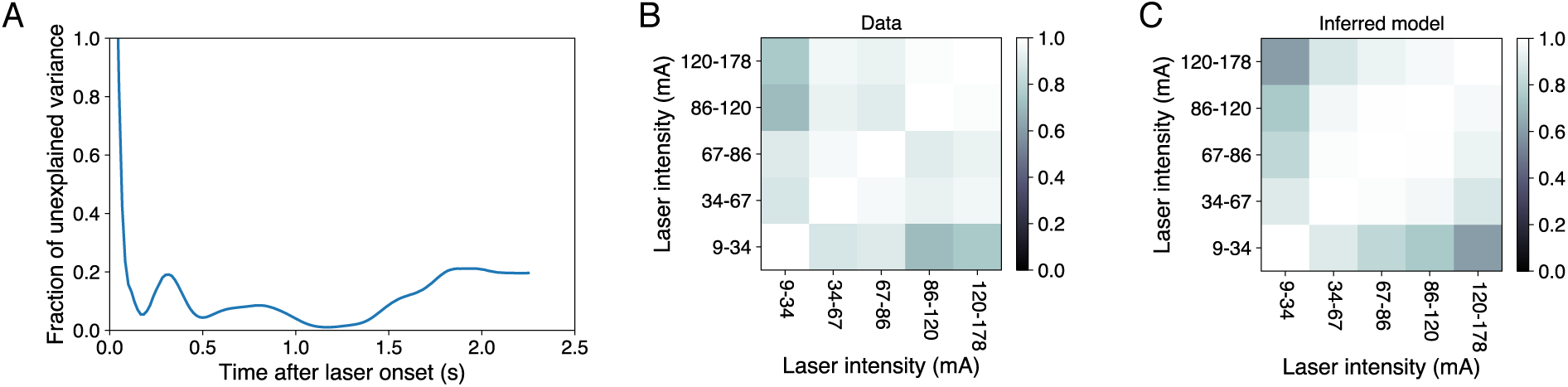
Comparison of the automated model to the manually curated model of Ref. [2]. (A) The fraction of the overall explainable velocity variance not explained by our model; the unexplained variance is roughly half of that of Ref. [2]. (B, C) The escape response has been modeled previously as stereotypical, with an overall scaling that is laser current dependent (for laser currents > 25 mA). Perfect stimulus-dependent rescaling of the escape would correspond to a correlation of 1 between velocity traces for different stimuli. (B) shows these correlations, over time, between mean response speeds in different laser current bins. (C) shows the same correlations in the inferred model. To the extent that correlations in both panels are nearly the same, our model recovers the approximate stereotypical nature of the response [2].

## Acknowledgments

This work was partially supported by NIH Grants EB022872 and NS084844, James S. McDonnell Foundation Grant 220020321, NSERC Discovery Grant, and NSF Grant IOS-1456912. We are grateful to the NVIDIA corporation for donated Tesla K40 GPUs. We are grateful to the Aspen Center for Physics, supported by NSF grant PHY-1607611, for supporting this research within “Physics of Behavior” programs.

## References

[1] A. Mohammadi, J. Byrne Rodgers, I. Kotera, and W. Ryu, BMC Neurosci 14, 66 (2013).

[2] K. Leung, A. Mohammadi, W. Ryu, and I. Nemenman, PLoS Comput Biol 12, e1005262 (2016).

[3] M. Schmidt and H. Lipson, Science 324, 81 (2009).

[4] M. Schmidt, R. Vallabhajosyula, J. Jenkins, J. Hood, A. Soni, J. Wikswo, and H. Lipson, Physical Biology 8, 055011 (2011).

[5] D. Sussillo and L. Abbott, Neuron 63, 544 (2009).

[6] G. Neuert, B. Munsky, R. Tan, L. Teytelman, M. Khammash, and A. van Oudenaarden, Science (New York, NY) 339, 584 (2013).

[7] B. Daniels and I. Nemenman, Plos One 10, e0119821 (2015).

[8] B. Daniels and I. Nemenman, Nature Commun 6, 8133 (2015).

[9] S. Brunton, J. Proctor, and J. Kutz, Proc Natl Acad Sci (USA) 113, 3932 (2016).

[10] N. Mangan, S. Brunton, J. Proctor, and J. Kutz, IEEE Transactions on Molecular, Biological and Multi-Scale Communications 2, 52 (2016).

[11] Z. Lu, J. Pathak, B. Hunt, M. Girvan, R. Brockett, and E. Ott, Chaos: An Interdisciplinary Journal of Nonlinear Science 27, 041102 (2017).

[12] J. Pathak, B. Hunt, M. Girvan, Z. Lu, and E. Ott, Phys Rev Lett 120, 024102 (2018).

[13] C. Pandarinath, D. O’Shea, J. Collins, R. Jozefowicz, S. Stavisky, J. Kao, E. Trautmann, M. Kaufman, S. Ryu, L. Hochberg, J. Henderson, K. Shenoy, L. Abbott, and D. Sussillo, bioRxiv, 152884 (2017).

[14] A. Henry, M. Hemery, and P. François, PLoS Comput Biol 14, e1006244 (2018).

[15] P. Nelson, Physical Models of Living Systems (W. H. Freeman and Co., New York, NY, 2015).

[16] V. Vapnik, Statistical learning theory (Wiley, 1998).

[17] J. Rissanen, Stochastic Complexity in Statistical Inquiry Theory (World Scientific, 1989).

[18] W. Bialek, I. Nemenman, and N. Tishby, Neural Computation 13, 2409 (2001).

[19] M. Savageau and E. Voit, Mathematical biosci 87, 83 (1987).

[20] R. Beer and B. Daniels, “Saturation probabilities of continuous-time sigmoidal networks,” (2010), arXiv: 1010.1714.

[21] D. J. MacKay, Neural Comput 4, 415 (1992).

[22] V. Balasubramanian, Neural Comput 9, 349 (1997).

[23] R. Eaton, Neural Mechanisms of Startle Behavior (Springer US, 2013).

[24] J. Pirri and M. Alkema, Curr Opin Neurobiol 22, 187 (2012).

[25] C. Bargmann, J. Thomas, and H. Horvitz, Cold Spring Harbor Symposia on Quantitative Biology 55, 529 (1990).

[26] M. Hilliard, A. Apicella, R. Kerr, H. Suzuki, P. Bazzicalupo, and W. Schafer, The EMBO Journal 24, 63 (2005).

[27] J. Kaplan and H. Horvitz, Proc Natl Acad Sci USA 90, 2227 (1993).

[28] N. Wittenburg and R. Baumeister, Proc Natl Acad Sci USA 96, 10477 (1999).

[29] G. J. Stephens, B. Johnson-Kerner, W. Bialek, and W. S. Ryu, PLoS computational biology 4, e1000028 (2008).

[30] M. B. Goodman and P. Sengupta, Pflugers Archiv : European journal of physiology 470, 839 (2018).

[31] S. Liu, E. Schulze, and R. Baumeister, PloS one 7, e32360 (2012).

[32] J. G. White, E. Southgate, J. N. Thomson, and S. Brenner, Philosophical transactions of the Royal Society of London. Series B, Biological sciences 314, 1 (1986).

[33] S. Gao, L. Xie, T. Kawano, M. D. Po, S. Guan, M. Zhen, J. K. Pirri, and M. J. Alkema, Nature communications 6, 6323 (2015).

[34] V. Sourjik and N. S. Wingreen, Current opinion in cell biology 24, 262 (2012).

[35] A. Paulick, V. Jakovljevic, S. Zhang, M. Erickstad, A. Groisman, Y. Meir, W. S. Ryu, N. S. Wingreen, and V. Sourjik, eLife 6 (2017), 10.7554/eLife.26607.

[36] A. Jilkine and L. Edelstein-Keshet, PLoS Comput Biol 7, e1001121 (2011).

[37] A. Levchenko and P. Iglesias, Biophys J 82, 50 (2002).

[38] P. Mehta, M. Bukov, C. Wang, A. Day, C. Richardson, C. Fisher, and D. Schwab, arXiv.org (2018), 1803.08823v1.

[39] M. Transtrum, B. Machta, K. Brown, B. Daniels, C. Myers, and J. Sethna, J Chem Phys 143, 010901 (2015).

[40] C. Myers, R. Gutenkunst, and J. Sethna, Computing in Sci Engin 9, 34 (2007).

[41] A. Mohammadi, Quantitative Behavioral Analysis of Thermal Nociception in Caenorhabditis Elegans: Investigation of Neural Substrates Spatially Mediating the Noxious Response, and the Effects of Pharmacological Perturbations, Ph.D. thesis, University of Toronto (2013).

